# Agonal factors distort gene-expression patterns in human postmortem brains

**DOI:** 10.1101/2020.07.11.198523

**Authors:** Jiacheng Dai, Yu Chen, Chao Chen, Chunyu Liu

## Abstract

Agonal factors, the conditions that occur just prior to death, can impact the molecular quality of postmortem brains, influencing gene expression results. Nevertheless, study designs using postmortem brain tissue rarely, if ever, account for these factors, and previous studies had not documented nor adjusted for agonal factors. Our study used gene expression data of 262 samples from ROSMAP with the following terminal states recorded for each donor: surgery, fever, infection, unconsciousness, difficulty breathing, and mechanical ventilation. Performed differential gene expression and weighted gene co-expression network analyses (WGCNA), fever and infection were the primary contributors to brain gene expression changes. Fever and infection also contributed to brain cell-type specific gene expression and cell proportion changes. Furthermore, the gene expression patterns implicated in fever and infection were unique to other agonal factors. We also found that previous studies of gene expression in postmortem brains were confounded by variables of hypoxia or oxygen level pathways. Therefore, correction for agonal factors through probabilistic estimation of expression residuals (PEER) or surrogate variable analysis (SVA) is recommended to control for unknown agonal factors. Our analyses revealed fever and infection contributing to gene expression changes in postmortem brains and emphasized the necessity of study designs that document and account for agonal factors.

## Introduction

Postmortem brain samples are widely used for human genetic studies, primarily for the study of neuropsychiatric disorders such as schizophrenia, bipolar disorders and Alzheimer’s disease^1^. Genetic mechanisms studied in postmortem brains from individuals with neuropsychiatric disorders such as transcriptome patterns reflect the genetic impacts of a lifetime of severe mental illness as well as countless environmental factors. Correcting for the extraneous environmental factors that influence gene expression in postmortem tissue^2^ is critical to accurate data analyses. One such category of environmental factors, agonal factors, describe the conditions that occur before death. While they represent an inevitable environmental component, agonal factors are historically neglected in postmortem brain studies.

Agonal factors refer to the manner of death and the terminal state before death. The manner of death include slow, intermediate, fast from natural causes, and violent fast^3^. Terminal states of consequence include coma, inadequate oxygen, fever, infection, and artificial respiration^4, 5^. Previous research has shown that messenger RNA (mRNA) is vulnerable to a greater or lesser degree to agonal factors^6, 7^. Some studies have provided an agonal factor score to account for specific agonal conditions and terminal states per individual^8, 9^. Agonal factors have also been reported to negative affect the gene expression profile^8 9^. Similar influences are induced by low tissue pH^10^. Such factors cause irreversible decomposition before and up to the moment of death. While some studies have incorporated agonal factors, the basic design typically retains the following shortcomings: 1) Studies focused primarily on the correlation of agonal factors to aspects such as the RNA integrity number, tissue pH, and whole gene expression pattern, neglect to identify the gene expression pathways that underlie agonal related regulation; 2) Such studies^8-11^ often roughly combine various terminal states together, thereby overlooking the potential interactions or conflicts between them; and, 3) Analysis if often limited to Pearson correlation and differential gene expression when further methods are needed to reveal the expanse of gene co-expression relationships.

We hypothesize that individual agonal factors uniquely alter postmortem brain gene expression and that successful adjustment of agonal-related variants within gene expression data is necessary and achievable. To investigate this matter, our study included 262 samples of postmortem human brain tissue from the Religious Orders Study and Memory and Aging Project (ROSMAP) study. The data included complete agonal information for the following durations for all samples: surgery, fever, infection, unconsciousness, difficulty breathing, and on mechanical ventilation. We performed differential gene expression and identified the gene co-expression network. We performed linear regression analysis to correct for agonal-related surrogate variables and hidden batch effects.

## Results

### Agonal factors are associated with gene expression in the human brain

Terminal state was recorded in the discovery data (ROSMAP) in our study. Details included *surgery, fever, infection, unconsciousness, difficulty breathing*, and *artificial ventilation*, which are recorded 3 days prior to death. In the following analysis, we focused primarily on these terminal states, performing differential gene expression analyses and gene co-expression analyses to uncover the potential effects that terminal states have on the gene regulatory network.

Differential gene expression (DEG) analysis showed that only fever, infection and unconsciousness were significantly associated with various gene expressions. We found 344 differentially expressed genes (DEGs) that were associated with *fever* (adjust p<0.05, figure 1A), 51 DEGs that were associated with infection (adjust p<0.05, figure 1B), and 5 DEGs that were associated with unconsciousness (adjust p<0.05). However, no significant DEGs were induced by *surgery, difficulty breathing* and artificial *ventilation*. These results showed that the agonal-related variants in data were mainly comprised of *fever* and *infection*, rather than *difficulty breathing* or *artificial ventilation*.

**Figure 1.**
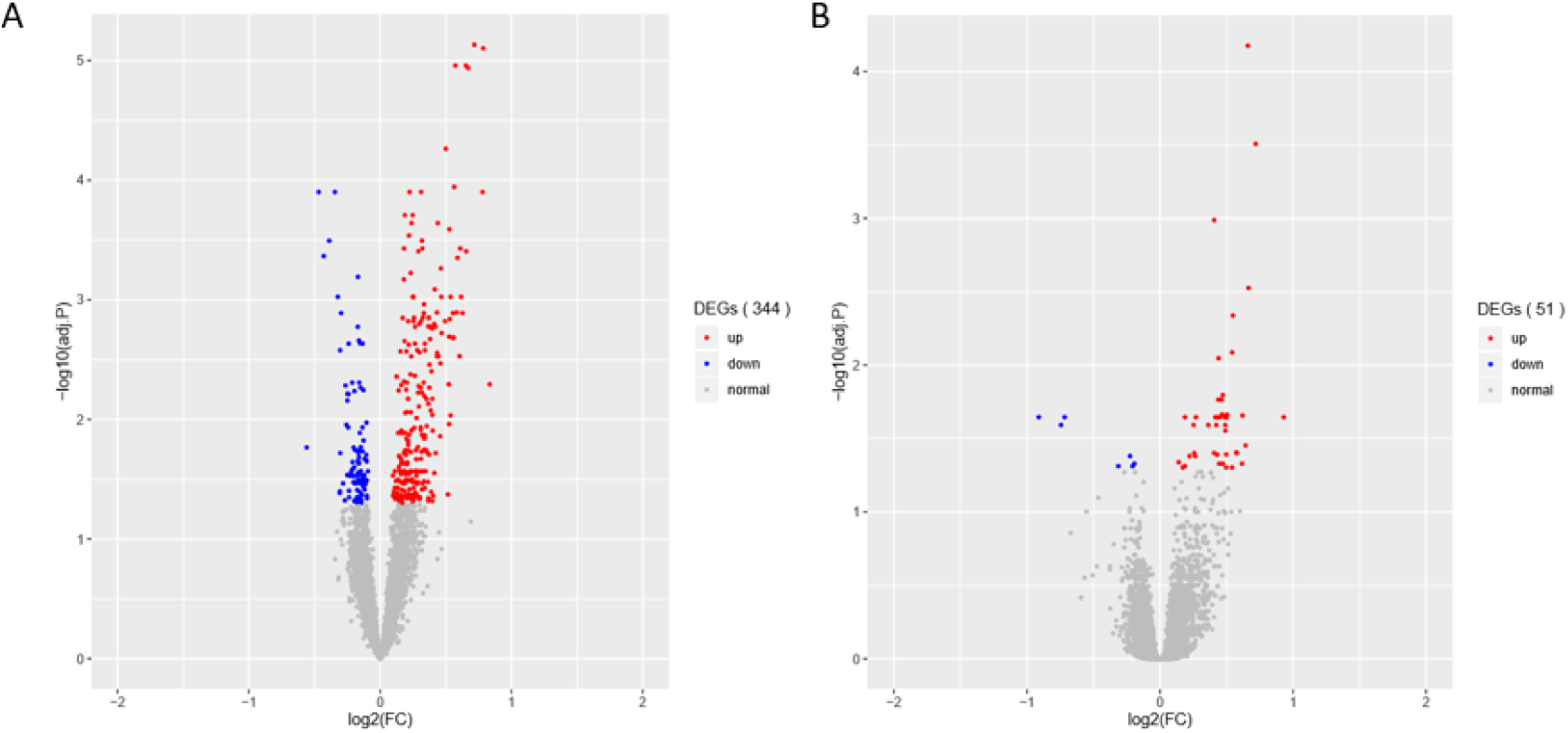
Differential expressed genes (DEGs) associated with the agonal factors of fever (a) and infection (b). In total, 344 significant DEGs were associated with agonal fever, and 51 significant DEGs were associated with infection.

The major variance components of bulk brain tissue may be driven by cell proportion, so we performed cell deconvolution to evaluate cell proportion in fever or infection. We found microglia (P=0.022) that showed significant differences in fever and non-fever sample. The cell proportion of microglia in fever sample (average 7.6%) is higher than in non-fever sample (average 5.9%). After we correct the cell proportion, we only found 11 fever-related DEGs and 1 infection-related DEGs. These results showed that fever may increase the cell proportion of microglia in pre-mortem brain.

We attempted to replicate the DEG results using the data from Hagenauer’s study (GSE92538). For AFS ranging from 0 to 3, we calculated DEGs of 0 versus 1, 0 versus 2, 0 versus 3. We found 1992, 2368, 2660 DEGs respectively in the replicated results. We also combined AFS of 1 to 3 together to compare with the AFS of 0, which resulted in 3529 DEGs. The replicated DEG results overlapped with fever-related DEGs with 147, 145, 211, and 247 genes (Table1). Fever-related DEGs are also significantly enriched in replicate DEG gene sets(Fisher p<0.05). However, there are few overlaps with *infection* DEGs (Table 1). The replicate results discovered same DEGs in *fever*, indicating that agonal factors’ effect did exist. However, we didn’t replicate the DEG results for *infection* due to the scarcity of recorded agonal data. Furthermore, the heterogeneity of criteria for the same agonal factor impedes proper replication.

**Table 1.** P value of cell proportion differential analysis.

|  | Ast | End | Ex | Mic | Oli | Per |
| --- | --- | --- | --- | --- | --- | --- |
| Fever | 0.926 | 0.936 | 0.36 | 0.022 | 0.936 | 0.06 |
| Infection | 0.36 | 0.936 | 0.36 | 0.36 | 0.98 | 0.753 |

**Table 2.** DEGs overlap of ROSMAP dataset and replicate dataset.

|  | DEGs of fever: 344 | DEGs of infection: 51 |
| --- | --- | --- |
| DEGs 0 vs 1: 1992 | 147 | 3 |
| DEGs 0 vs 2: 2368 | 145 | 1 |
| DEGs 0 vs 3: 2660 | 211 | 3 |
| DEGs 0 vs 1+2+3 : 3529 | 247 | 4 |

DEG gene ontology of *fever* and *infection* showed significant enrichment in synapse- and immune-related pathways (Figure 2). In the top gene ontology enrichment of DEGs for *fever* (Figure 2A), we identified genes strongly enriched for immune-related pathways. In addition, we found gene enrichment for other pathways, including the response to unfolded protein pathway (adjust p=0.00087), a protective response induced during periods of cellular stress that aims to restore protein homeostasis^12^. In the top gene ontology enrichment of DEGs for infection (Figure 2B), the strongest enrichment was that of the synapse organization pathway (adjust p=0.018831), followed by the cell killing (adjust p=0.018831), the regulation of synapse organization (adjust p=0.01983) and the regulation of synapse structure or activity (adjust p=0.01983) pathways, indicating that in addition to immune-related pathways, synapse-related gene expression pathways also play an important role in the brain during *infection*.

**Figure 2.**
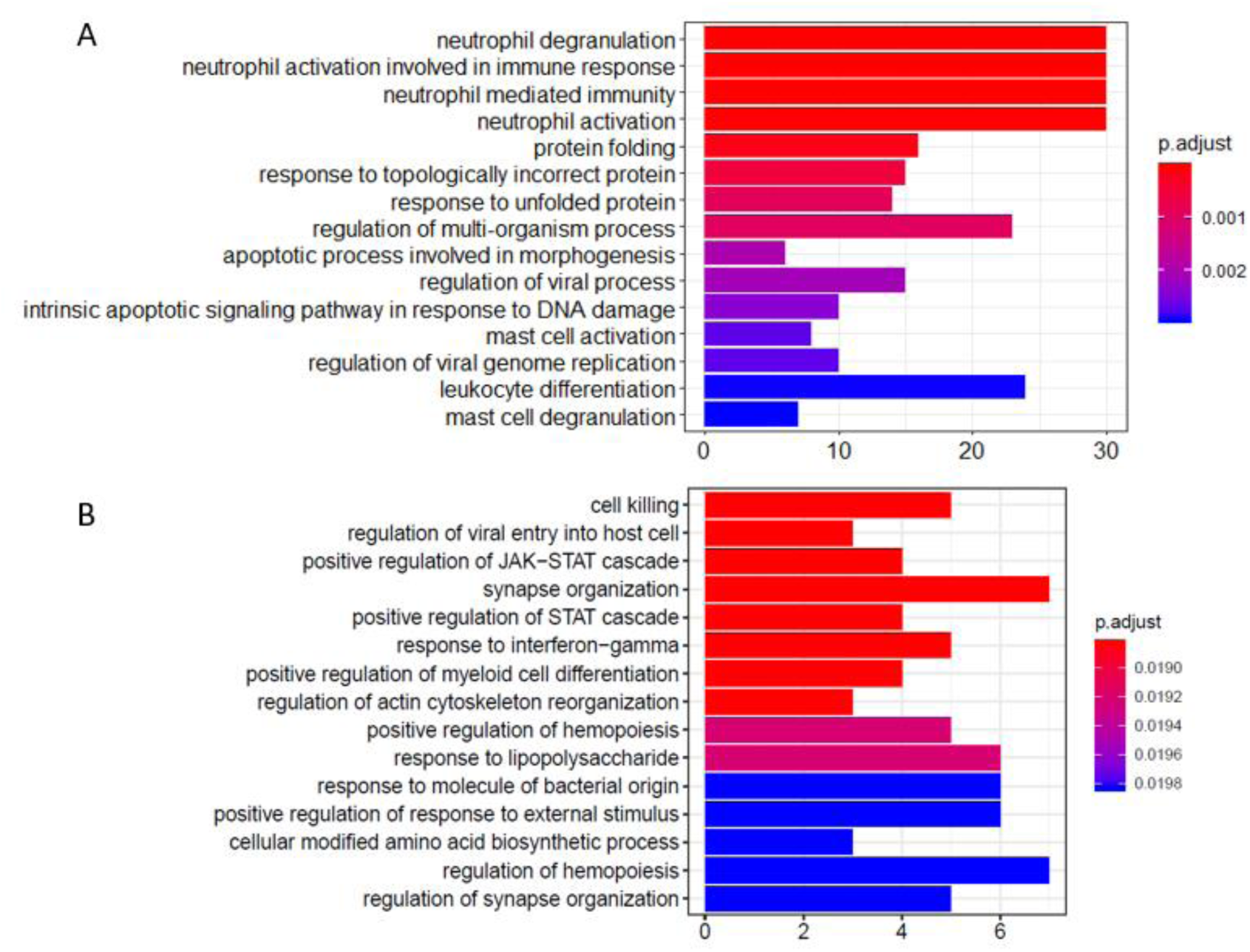
Gene ontology enrichment analysis of differentially expressed genes (DEGs) associated with fever (a) and infection (b). We found that DEGs associated with fever are enriched in immune- and apoptotic-related pathways, whereas DEGs associated with infection are enriched within synapse-related pathways.

### Correlation between terminal states

The correlation of log2-FC effect sizes across terminal states also show contrasting directional effects on gene expression related to terminal state (Figure 3). We found that *fever* positively correlated with *infection* and *unconsciousness* (*ρ*=0.49, 0.42); whereas, *difficulty breathing* positively correlated with *artificial ventilation* (*ρ*=0.37). We also observed that *fever* negatively correlated with *artificial ventilation* (*ρ*=-0.48), *infection* negatively correlated with *surgery* (*ρ*=- 0.44), and *unconsciousness* negatively correlated with *difficulty breathing* and *artificial ventilation* (*ρ*=-0.31, −0.38). Based on the log2-FC correlation, we concluded that the relationships among terminal states are complex and may have an opposite effect on gene expression.

**Figure 3.**
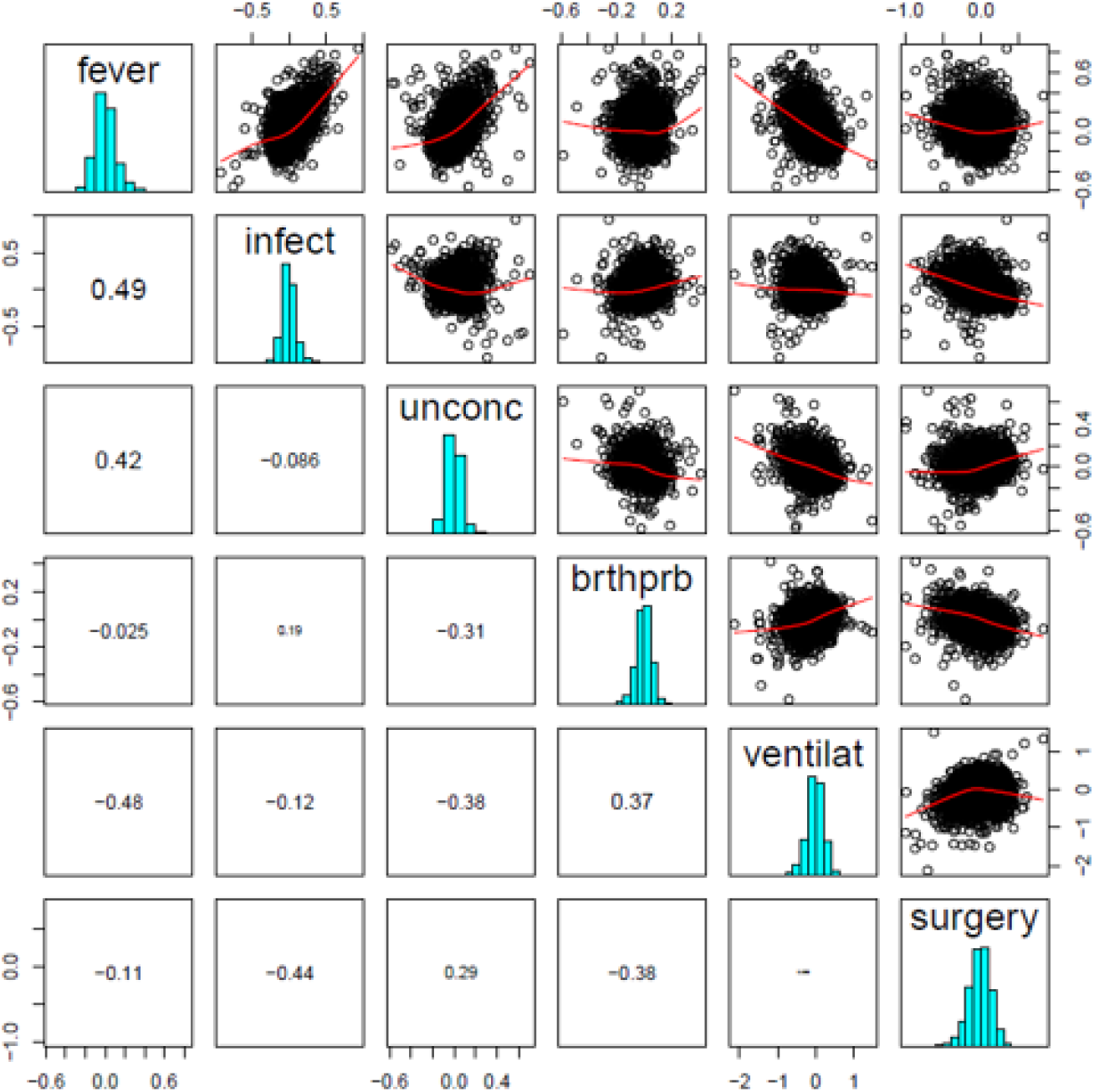
Terminal state associated differentially expressed gene (DEG) log2-FC effect size correlation. We found a significant positive correlation between fever and infection, difficulty breathing and artificial ventilation. We also a found negative correlation between fever and ventilation, infection and surgery, and unconsciousness and ventilation.

Considering that the correlation of effect size was caused by the correlation of clinical phenotypes, we checked whether terminal states correlated clinically to participants. We found that *fever* and *infection* were significantly correlated (P=0.0002, Fisher’s exact test), and that *fever* and *difficulty breathing* were significantly correlated (P=0.01, Fisher’s exact test). Only *fever* and *infection* showed significant correlation both in gene expression level and clinical level, indicating a close relationship within this data. Other terminal states which have correlation in gene expression level showed insignificant correlation clinically. These results indicated that the correlation of gene expression in different terminal state was not decided by clinical correlation.

### Agonal-related co-expression networks reveal cell type-related modules

Investigating a possible relationship between gene co-expression and agonal factors, we performed weighted correlation network analysis (WGCNA) on constructed gene co-expression networks (Figure 4A). We further analyzed whether gene co-expression modules (ME) using the following methods: cell-type enrichment analysis, psychiatric disorders’ candidate gene enrichment analysis and gene ontology analysis. Of the 18 identified modules, two upregulated modules (ME11, ME12) were significantly associated with the combination of *fever* and *infection* (FDR p<0.05); the ME8 was significantly associated with *fever*, while the ME17 was significantly associated with *infection*; whereas, only one downregulated module (ME10) was associated with *fever*. Yet, no modules were significantly associated with *surgery, unconscious*ness, *difficulty breathing* or *artificial ventilation*.

**Figure 4.**
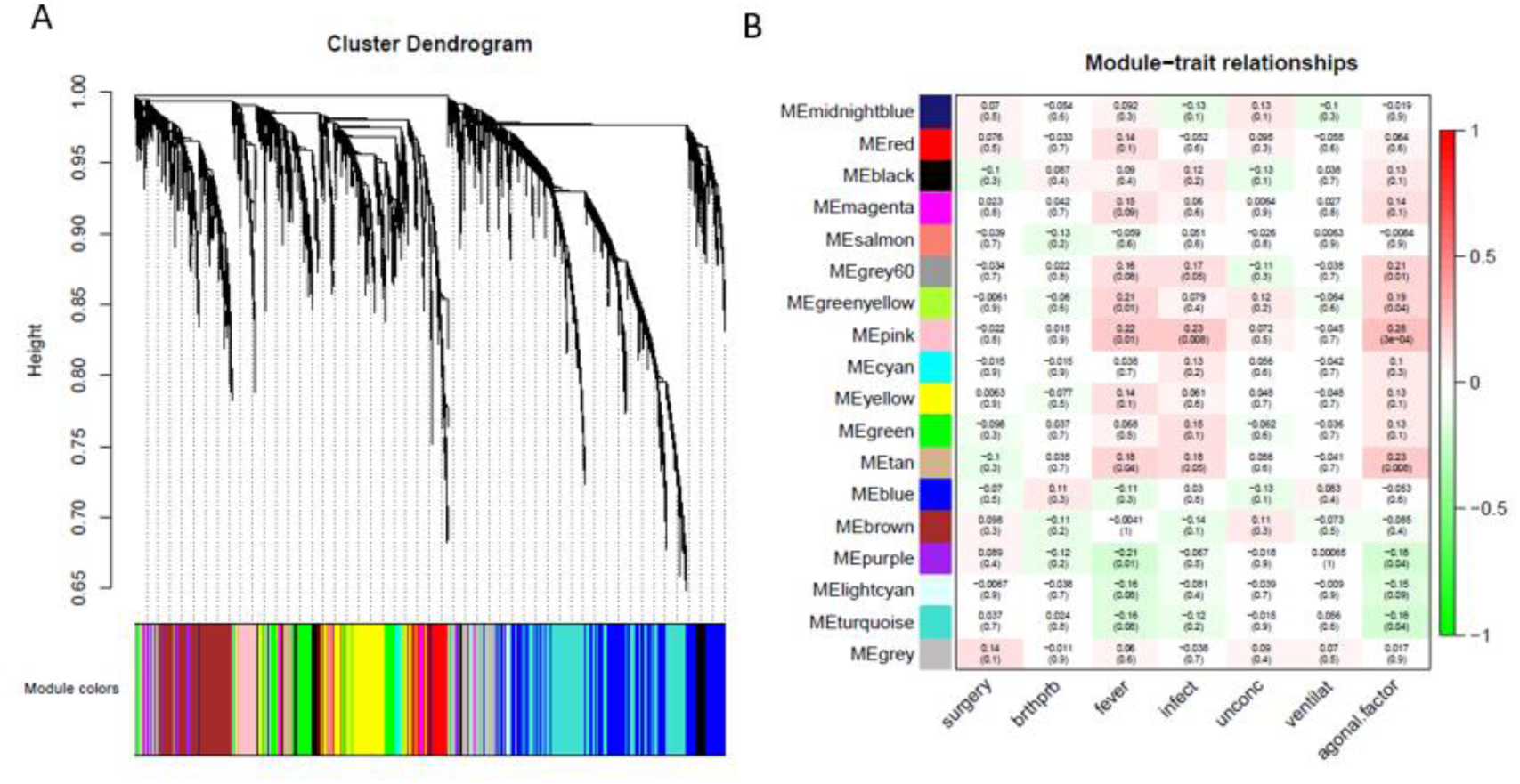
A network dendrogram (A) driven by a weighted gene co-expression analysis (WGCNA) and a module-trait correlation (B). In total, we identified 18 modules from the ROSMAP dataset. We also identified modules that were significantly correlated with agonal risk factors and the agonal factor score (AFS).

Correlating the gene co-expression module and terminal states shows us that terminal states introduce different effects on brain gene co-expression patterns (Figure 4B). Since *fever* and *infection* are frequently comorbid conditions in the clinical setting, we calculated AFS using *fever* and *infection* as a combined phenotype in the ROSMAP data for more significant results. We found five modules (ME17, ME8, ME11, ME12, ME10 and ME1) associated with the AFS calculated in this way. The ME1 subthreshold in the correlation analysis of *fever* and *infection* separated but reached significance in the correlation analysis of AFS.

An analysis of cell-type enrichment of the gene co-expression module revealed brain cell type-specific modules (Figure 5). The ME12 is enriched for microglia and endothelial cell types, the ME11 is enriched for microglia cell types, and the ME1 is enriched for neuron cell types. Different cell type played different role in brain function, so it is surprising that the upregulated modules are enriched for microglia and endothelial cell types, while the downregulated modules are enriched for neurons. The cell-type enrichment analysis indicates that agonal factors may have a cell type-specific impact on gene expression.

**Figure 5.**
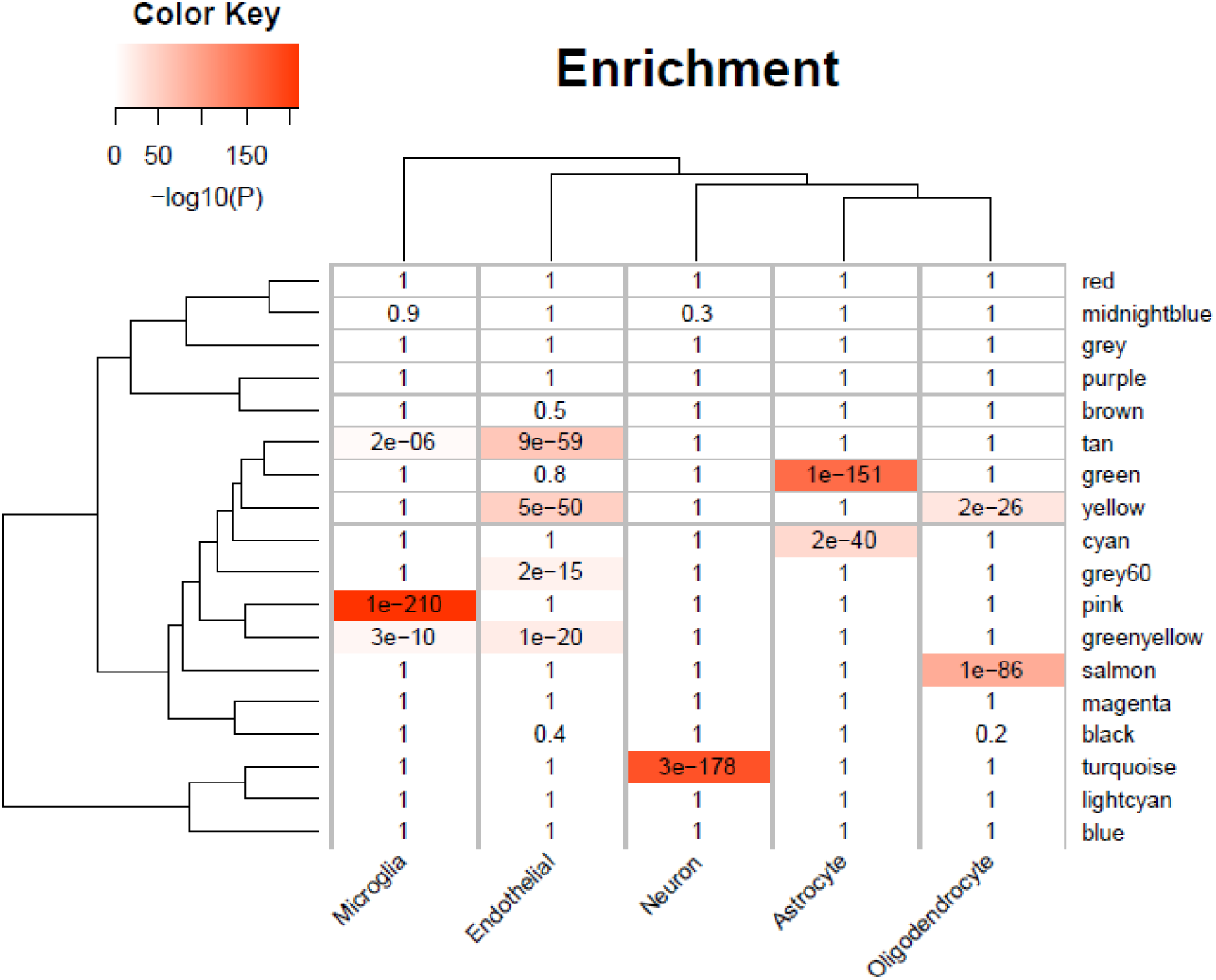
Cell-type specificity enrichment. We identified agonal-related modules enriched within neurons, microglia, and endothelial cells.

To test for module preservation of the ROSMAP data, we used data from Hagenauer et al.^9^ (GSE92538), whose group first proposed the concept of the AFS. For the modules which were associated with terminal states and AFS in the gene co-expression network of ROSMAP data, the ME8, ME11 and ME1 had Zsummary>10, which indicated that these modules were highly preserved in two datasets (Figure 6A). The ME12 show at least moderate evidence of preservation (Zsummary>5) in two datasets. The module preservation test indicated that several agonal-associated modules were conserved in different datasets. In the gene co-expression network of replicate data, we also observed similar cell-type enrichment of up- and down-regulated modules (Figure 6B, C), for down-regulated modules enriched for neuron and up-regulated modules enriched for microglia and endothelial cell types.

**Figure 6.**
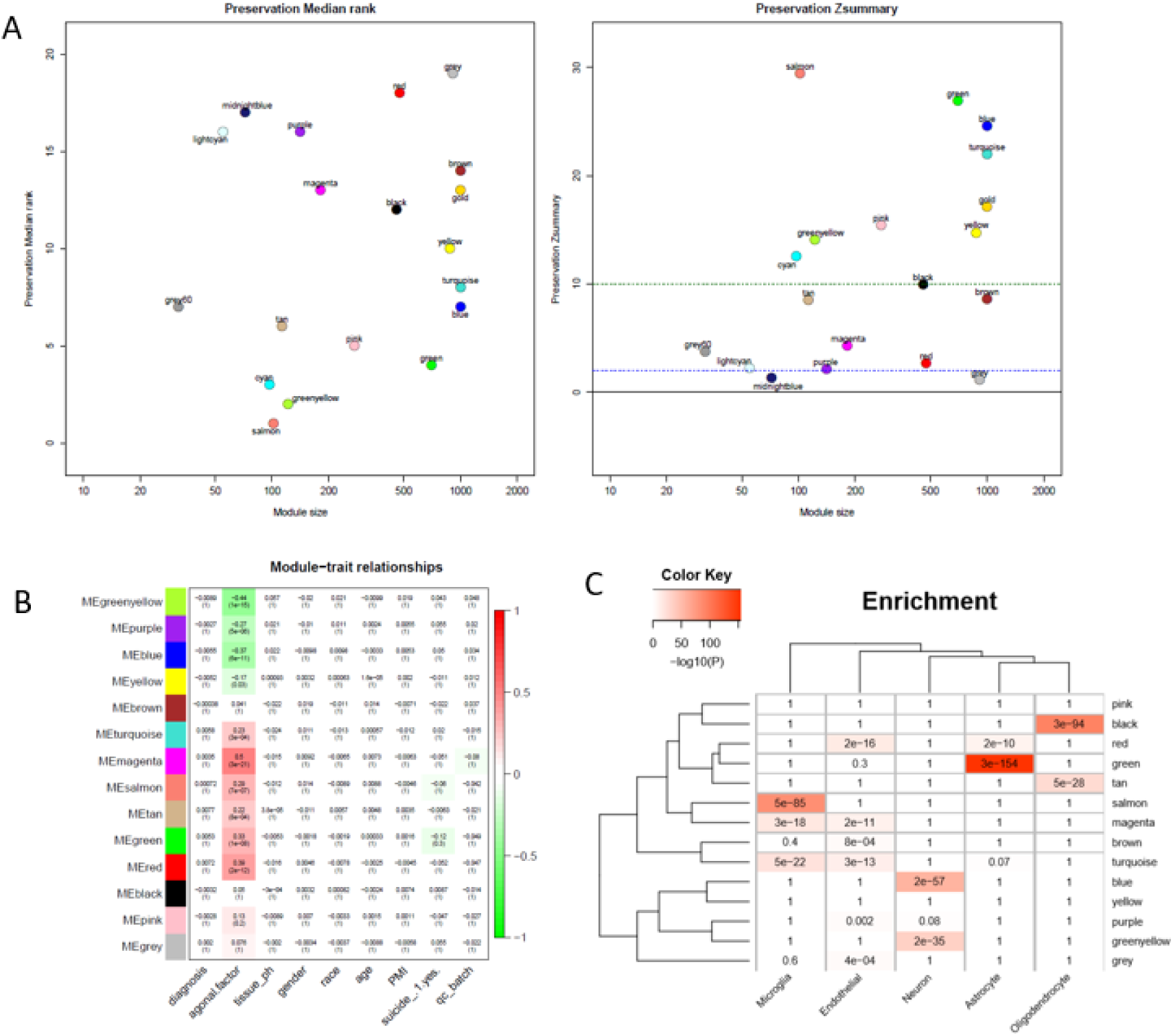
Replicated results using data from Hagenauer et al. (GSE92538). A. Module preservation test shows that modules from the ROSMAP dataset were conserved across different datasets; B. Module-trait correlation shows modules significantly correlated with the agonal factor score (AFS) in the replicate data; C. Cell-type specificity enrichment result shows agonal-related modules enriched within neurons and microglia in the replicate data.

We selected five cell type-enriched agonal-related gene co-expression modules (ME1, ME8, ME11, ME12, and ME17) as well as the fever/infection-related DEG to perform an analysis of psychiatric-disorder candidate gene enrichment. We used candidate genes of autism spectrum disorder^13-20^ (ASD), major depression disorder^16^ (MDD), and schizophrenia^13, 16, 17, 21-28^ (SCZ) from previous studies. The results showed the ME1, ME8, ME11 and ME12 significantly enriched in ASD and SCZ candidate genes (Figure 7). Also, we discovered that DEGs associated with fever were enriched in ASD and SCZ candidate genes which were discovered by co-expression network analysis.

**Figure 7.**
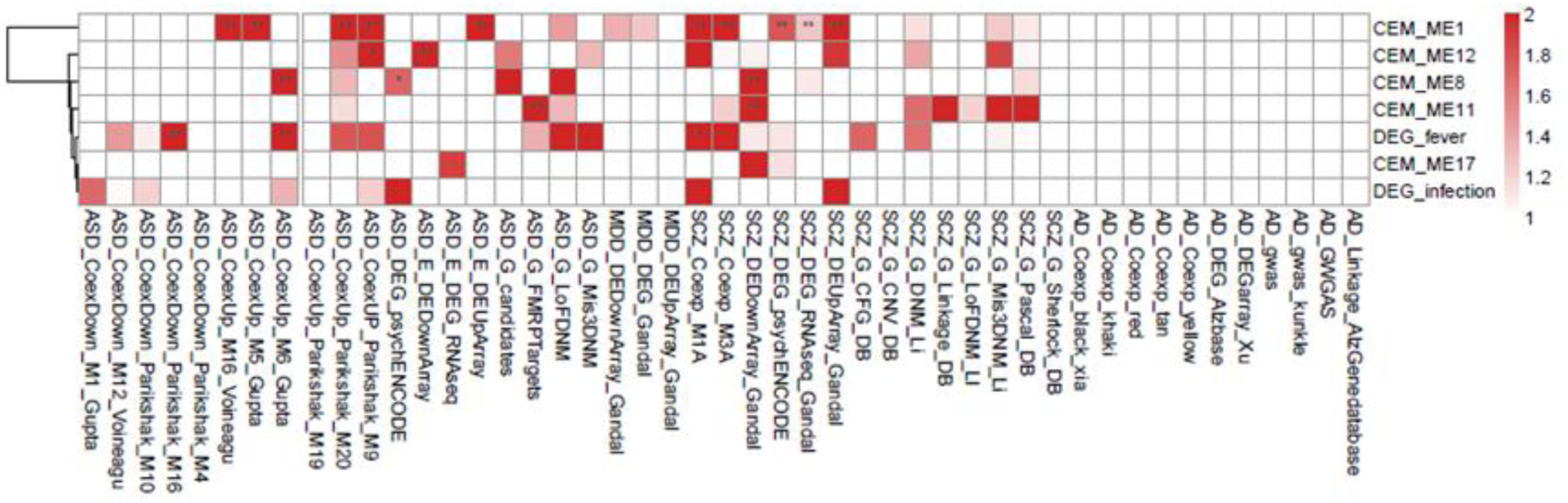
Gene enrichment analysis of the co-expressed module (CEM) genes and differentially expressed genes (DEGs) by psychiatric disorders type. We found that the schizophrenia (SCZ) and autism spectrum disorder (ASD) candidate genes previously identified in other studies are enriched in agonal-related modules.

The module shows a strong gene enrichment by cell-type, which can be associated with various biological processes. The ME1 was enriched for cell activity pathways, such as multicellular organismal homeostasis (adjust p=2.27E-05), response to extracellular stimulus (adjust p=2.51E-06), regulation of DNA-binding transcription factor activity (adjust p=4.23E-07), and immune-related pathways including T cell activation (adjust p=0.00295) (Figure 8A). The gene expression level of ME1 was down-regulated, suggesting that related gene expression pathway was suppressed. The ME12 strongly enriched for pathways such as the mRNA catabolic process, RNA catabolic process, translational initiation, and establishment of protein localization to organelle (Figure 8B). The gene expression level of the ME12 was up-regulated, which means that these cellular functions are activated to rescue the cell’s basic function. The ME11, which was enriched in microglia and endothelial cells, represents the biological processes of glial cells (Figure 8C). The top gene ontology enrichment pathways of this module were synapse-related, including pre-synaptic, endomembrane system organization, modulation of chemical synaptic transmission and synapse organization. The up-regulating of synapse-related function indicated that microglia and endothelial cells are activated to protect neuron cells during the terminal state.

**Figure 8.**
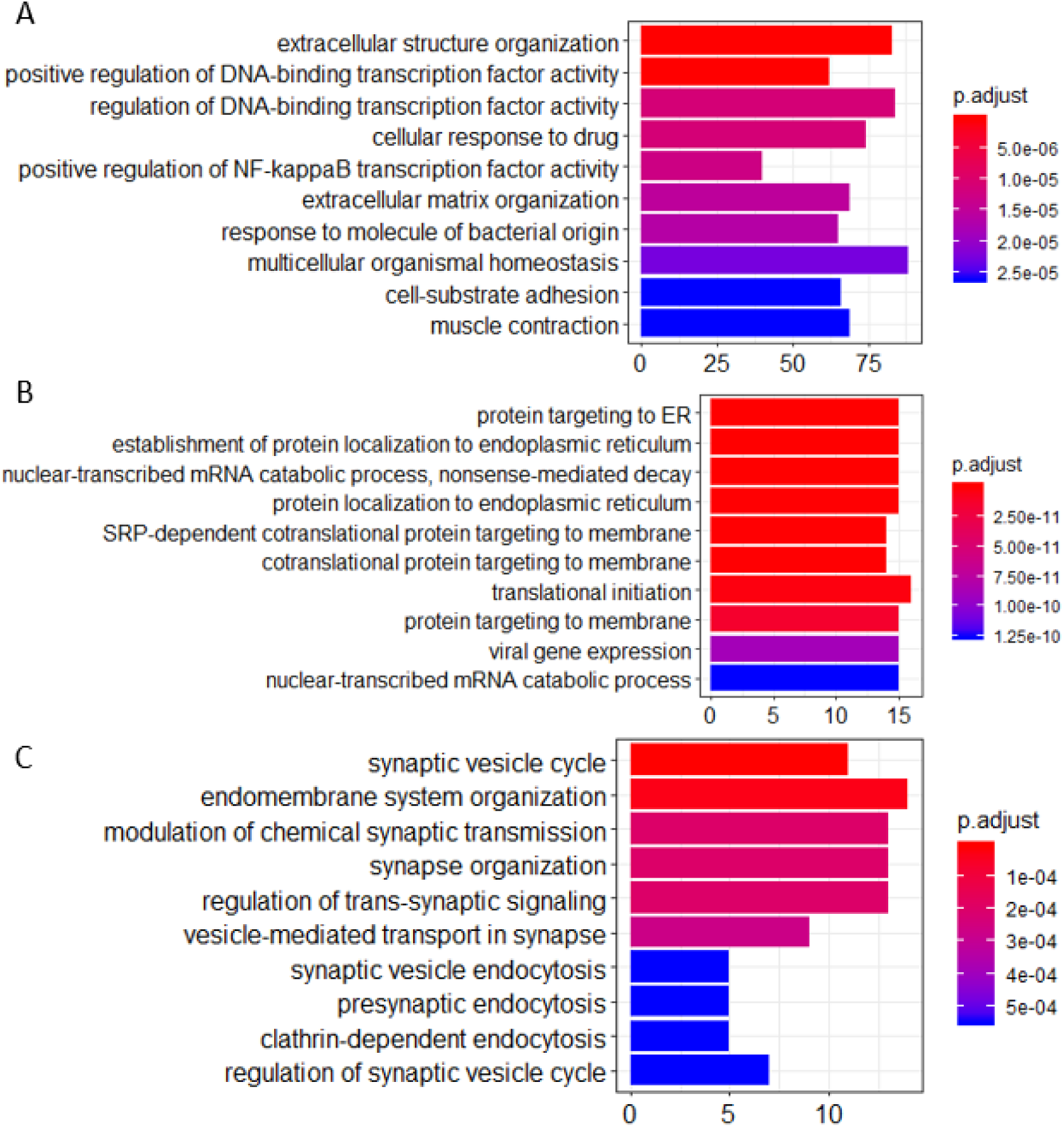
Gene ontology enrichment analysis of agonal-related modules: A. ME1; B. ME12; and C. ME11.

### Utilizing unknown factors analysis to predict agonal factors

In modern, high-throughput biomolecular experiments, unmeasured or unmodeled factors can confound the primary variables and confuse the results. Researchers usually use a hidden factors estimation method to model large-scale noise dependence. This dependence can be caused by unmeasured or unmodeled factors. These models include surrogate variable analysis (SVA) for gene expression data and PEER, designed for transcriptomic data from eQTL analysis. In our study, we used SVA and PEER to detect and correct for agonal factors. This also assisted in simulating agonal factors that were not otherwise recorded. For SVA, we detected 16 surrogate variables and correlated the 16 surrogate variables (sv) with terminal states (Figure 9A). Results showed that 10 of the 16 surrogate variables are significantly correlated with at least one agonal factor. We also observed that several surrogate variables correlated with more than 3 agonal factors. For example, sv2 is positively correlated with *surgery* but negatively correlated with *infection* and AFS; while, sv10 is negatively correlated with breathing difficulty, *fever, infection* and the AFS. These results also suggest a reverse effect for surgery and infection and a similar effect for *fever* and *infection*. In PEER analysis, we detected 15 hidden factors that explained the variants in the data (Figure 9B). For 15 hidden factors, we found 10 factors significantly correlated with terminal states. Similarly to surrogate variable correlation, *surgery* and *infection* showed an opposite correlation to hidden factor 2; while *fever, infection*, and the AFS showed correlation in the same direction for hidden factors 4 and 13.

Linear regression analysis succeeded in correcting for agonal-related surrogate variables (SVA) and for hidden factors (PEER). We performed principal variance component analysis (PVCA) of the gene expression matrix before and after correcting for 10 agonal-related surrogate variables (sv 1, 2, 4, 5, 6, 7, 9, 10, 12, and 14). Results showed a decreased variance for most phenotypes (Figure 11A, B). After selecting race as a phenotype of interest, we found that the variance of race increased after correction in SVA. The variance of mechanical ventilation also increased, due to a lack of its correlation to surrogate variables. We also performed PVCA on the quantile normalized gene expression matrix before and after correcting for the 10 agonal-related hidden factors (factors 1, 2, 4, 5, 8, 9, 10, 12, 13, and 15). Terminal state variation decreased except for the factor of mechanical ventilation, due to the lack of its correlation also with hidden factors (Figure 11C, D). After performing differential gene expression analysis following correction for hidden factors in PEER and surrogate variables in SVA, we found no DEGs for *fever* nor for *infection*. In conclusion, if researchers have not collected agonal related phenotypes, correction for unknown factors can still occur using such methods as SVA or PEER.

**Figure 10.**
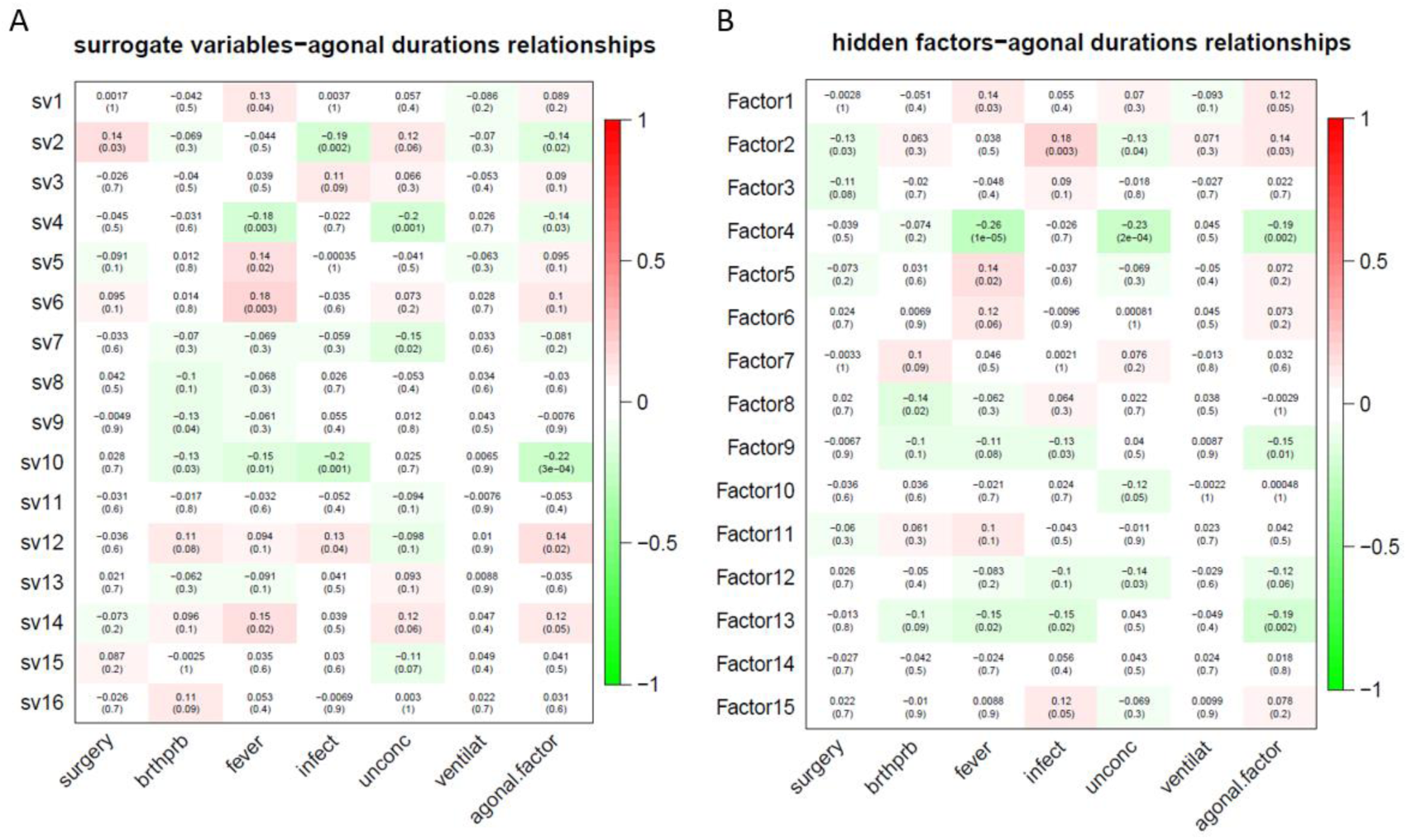
Unknown factors discovered by SVA (A) and PEER (b) and terminal state correlation. A. Surrogate variables 1, 2, 4, 5, 6, 7, 9, 10, 12, and 14 have a significant correlation with terminal states; B. Hidden factors 1, 2, 4, 5, 8, 9, 10, 12, 13, and 15 have a significant correlation with terminal states.

**Figure 11.**
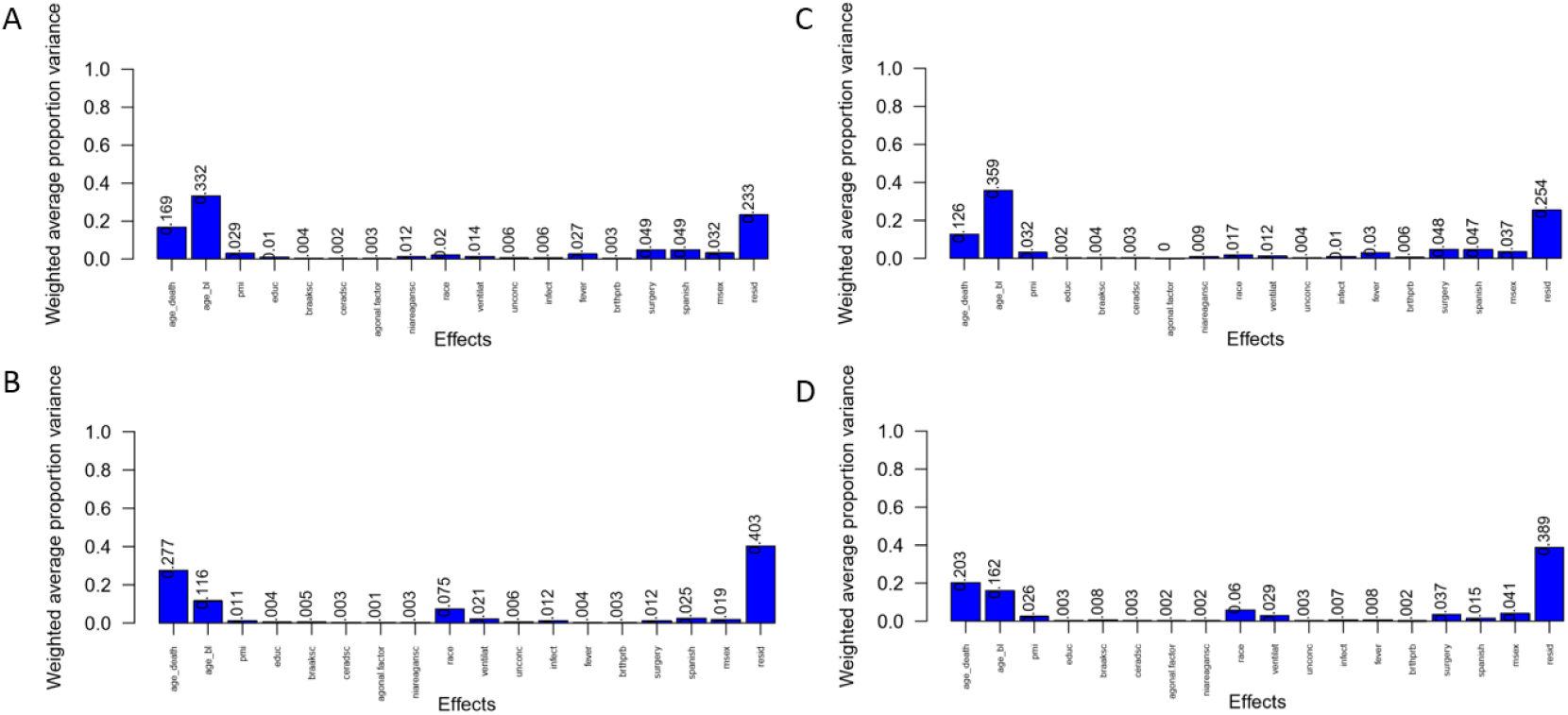
PVCA plot of correlation before and after correction of unknown variables. A. PVCA plot before the correction of surrogate variables; B. PVCA plot after the correction of surrogate variables; C. PVCA plot before the correction of hidden factors; D. PVCA plot after the correction of hidden factors.

We investigated previous studies’ results and found them enriched in genes associated with the agonal-related module. We then applied SVA to these datasets and evaluated the variables from agonal factors. We used microarray data of SCZ, ASD and IBD, and we compared the DEGs before and after the SVA adjustment. Performed a meta-analysis of 5 SCZ microarray data, we found 2044 DEGs of SCZ (FDR<0.05). After correcting 28 surrogate variables which had no significant correlation by disease group, we found 474 DEGs with 1628 genes filtered. Filtered genes were overlapped with fever-related DEGs (63 overlapped genes) and enriched in the ME8 (p=0.006) and the ME12 (p=0.004). Filtered genes were also enriched in hypoxia and oxygen level related pathways (Figure 12). We also found filtered genes in ASD as well as filtered genes enriched in oxygen levels-related pathways (Figure 13). However, IBD filtered genes were not enriched in agonal-related pathways (Figure 13).

**Figure 12.**
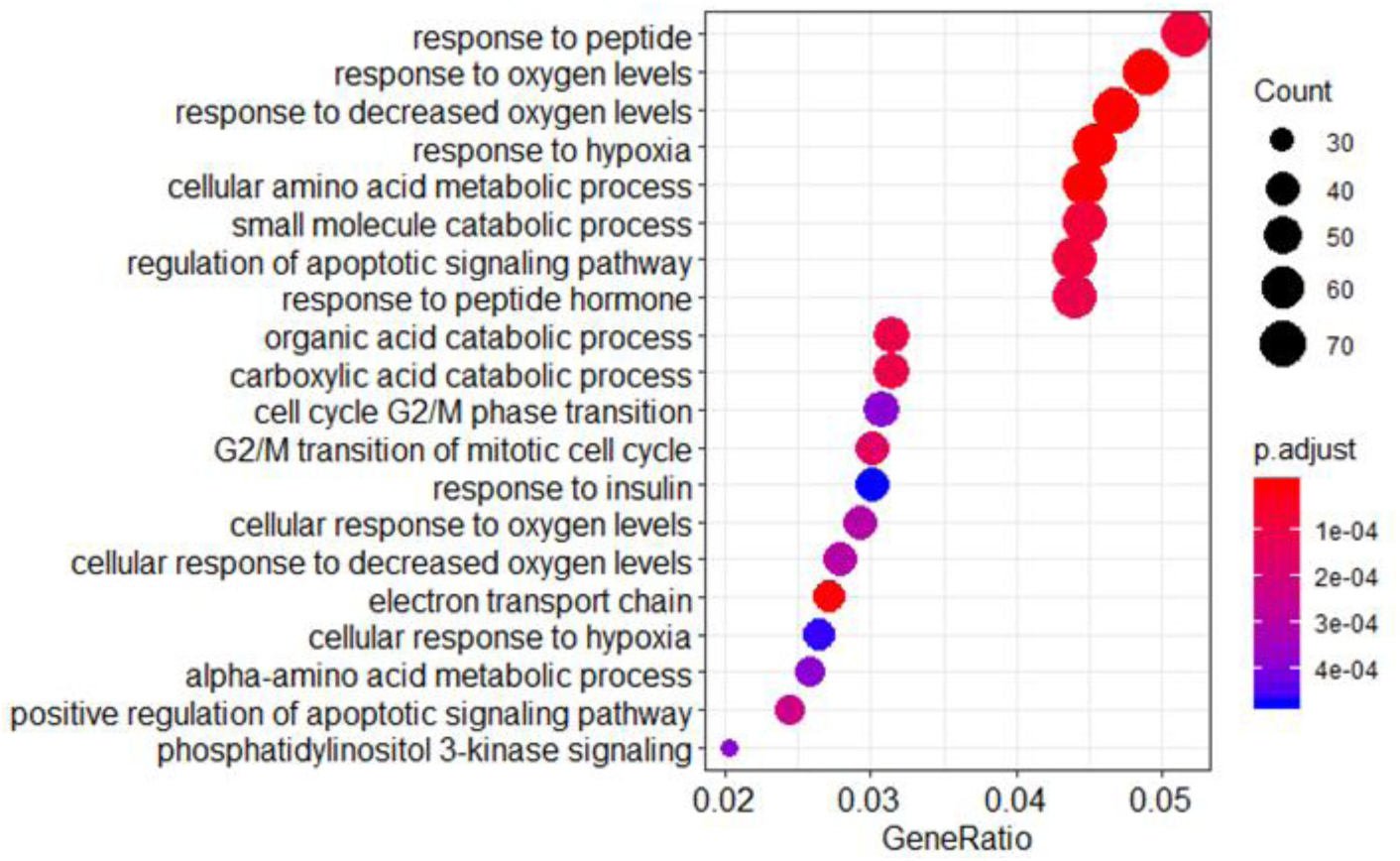
Gene ontology enrichment analysis of SCZ filtered genes.

**Figure 13.**
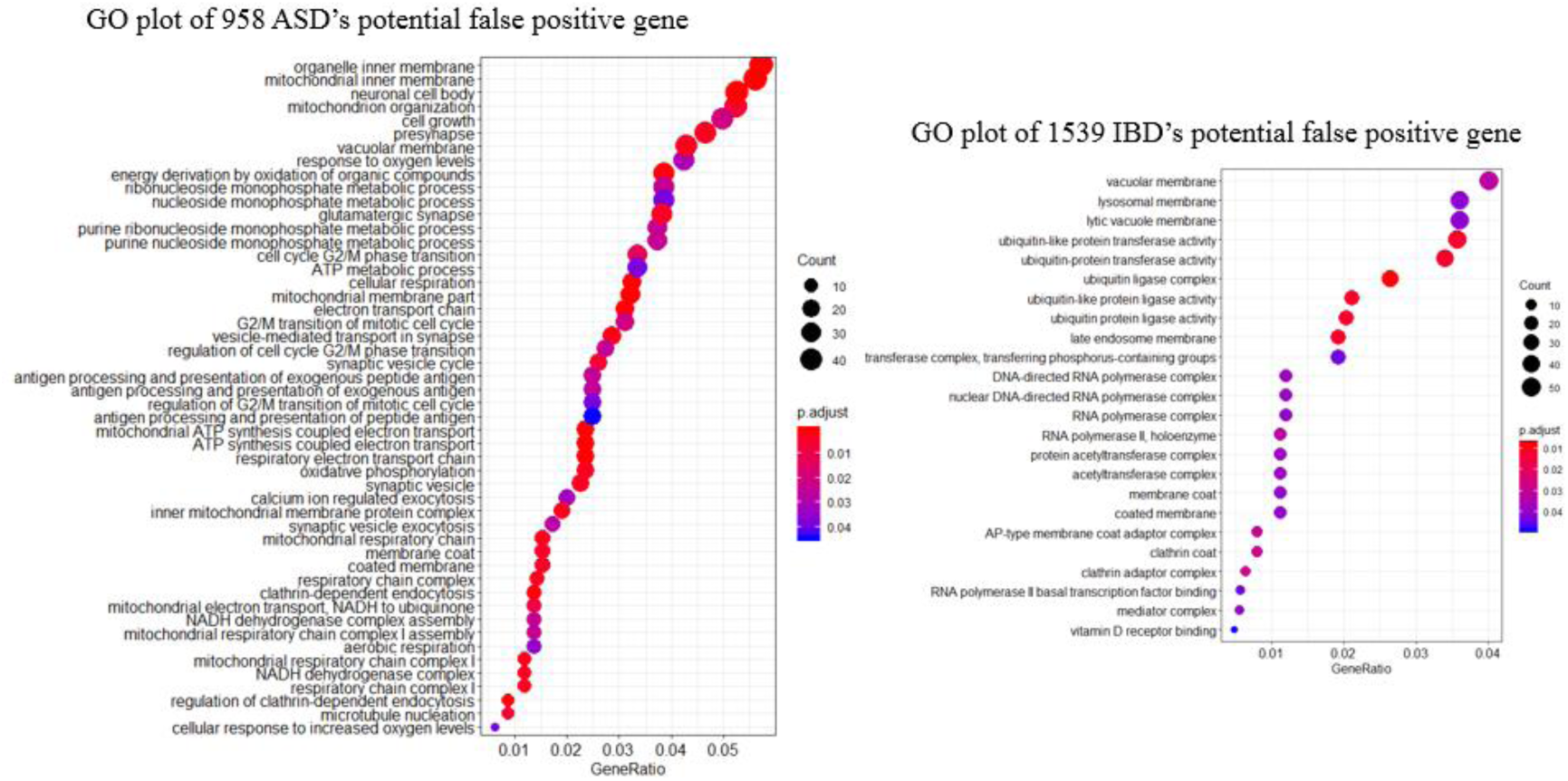
Gene ontology enrichment analysis of those genes related to autism spectrum disorder and irritable bowel disorder.

**Table 2.**
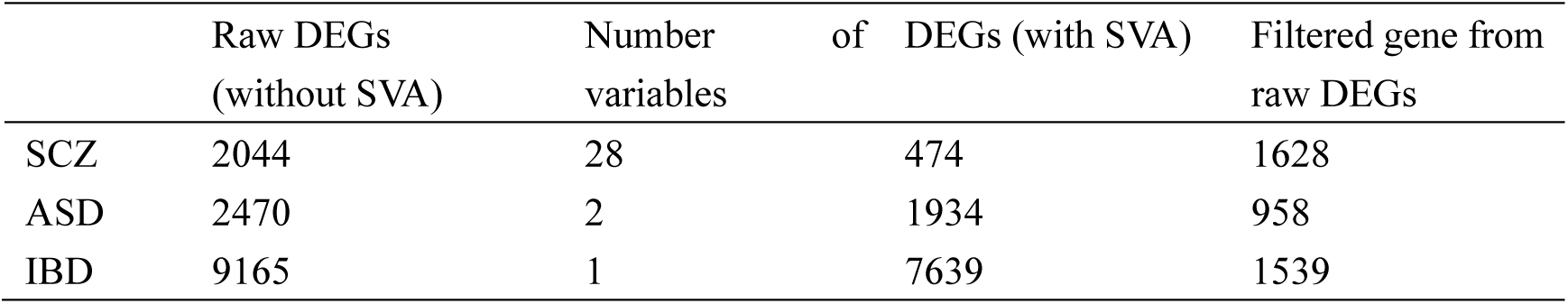
Data replication for SVA correction.

## Discussion

We present a transcriptomic data analysis of human postmortem brain from a public database, which provides a framework for understanding how different terminal states contribute to gene expression changes in postmortem brain tissue. We performed data analysis to identify genes that are expressed differently in various abnormal agonal conditions compare to normal post-mortem controls. Many genes show altered expression after undergoing an agonal state characterized mainly by fever and infection, enriched for cellular stress-related pathways. Cell proportion of microglia and neuron also showed alteration in fever and infection samples. To determine whether agonal factors related to other established disease candidate genes and to annotate its functional role in the human brain, we generated agonal-related co-expression networks. We identified Gene co-expression modules that significantly correlate with combination of fever and infection, drawing in positively co-expressed oligodendrocyte and microglia cell types as well as negatively co-expressed neuron cell types strongly enriched for SCZ and ASD candidate genes. Being a biological factor, agonal factors can be adjusted by linear regression analysis or unknown factors adjustment methods. Surprisingly, we found that data from previous studies may be confounded by agonal factors that were not accounted for. We found gene sets from studies results were enriched for hypoxia- and oxygen level-related pathways, while the gene sets can be filtered by SVA. Our study emphasizes that agonal factors are important biological factors, and that agonal factors should be documented and adjusted in postmortem brain tissue studies.

We performed data analysis aimed at fever, infection, unconsciousness, breathing difficulty, artificial respiration and premortem surgery, only fever and infection related to altered gene expression levels during the agonal period. We hypothesis the alteration possibly reflected by brain cell proportion changes^1^. According to cell deconvolution, we found cell proportion of microglia rises in fever sample. Similar pattern was found in gene co-expression network. We found fever- and infection-related modules are enriched for microglia-specific cell markers, which showed up-regulation for gene expression. The coincident results of cell type changes indicate that specific brain cell types have different sensitivity to terminal states. In fever, parts of the brain becomes inflamed, which may cause microglia high expressed as active immune defense. Besides, gene co-expression modules are also enriched for neuron- and endothelial-specific cell markers. The results indicated that, in stressed brain environment of agonal, neuron may be more vulnerable to the agonal state as compared with microglia and endothelial cells. One previous study reported that hypoxia was associated with increasing endothelial-specific expression and decreasing neuron-specific expression^9^. The combination of fever and infection may also be associated with brain hypoxia, reflecting the vulnerability of neurons to low oxygen and/or severe infection^29^. Besides, we found fever may activate an cellular response to unfolded protein pathways after exposure to a stressed environment. It was reported that in cancer cells, hypoxia can activate components of this pathway^30^. Agonal factors other than fever and infection did not show any significant DEGs nor any associated modules within the dataset. We compared the log2-FC of all agonal factors and found a negative correlation between artificial respiration and fever. Likewise, our module-trait correlation analysis yielded similar phenomena. When we combined all agonal factors to calculate AFS (i.e., surgery, difficulty breathing, fever, infection, unconsciousness, and artificial respiration), no modules correlated significantly to AFS. However, when we calculated AFS based on fever and infection, a greater number of gene co-expression modules showed significant correlation with AFS. Previous studies simply added the manner of death with terminal states to define the severity of agonal conditions, but our findings suggest that that their method may confound the various agonal effects. We found that terminal states may contribute uniquely to gene expression. Therefore, we hypothesize that differences of gene expression come from the consequences brought on by the various agonal factors. While some agonal factors, such as medical interventions like artificial respiration try to return the brain’s extracellular environment to normal, other conditions, such as fever and infection, intensify stress upon the brain’s extracellular environment.

Some researchers have addressed the concern that agonal factors may represent considerable confounders. For example, the Netherland Brain Bank suggested that it is necessity of recording agonal factors; furthermore, they emphasize that researchers should ensure that patient and control groups match for as many known confounding factors as possible, including agonal states and stress of dying^31^. In another example, Ramaker et al. sought to avoid the variability of agonal factors connected to an extended dying process. They included only post-mortem brain samples from individuals who experienced violent fast deaths ^32^. Nevertheless, most postmortem brain studies still did not account for agonal confounders. We checked the expression data from the public database BrainEXP^33^ and found that most of the contributing datasets did not collect agonal information. This is also the reason that we were unable to find another comprehensive dataset to replicate results. Moreover, studies proved to have an inconsistent definitions to the various agonal states, making result replication problematic. Datasets with larger sample sizes are needed in addition to comprehensive recordings of agonal factors in order to accurately evaluate their effects.

Previous gene expression analysis of post-mortem brains may be confounded by terminal states, which may have resulted in data errors and false positive results. Our agonal-related modules revealed the enrichment of several candidate genes from previous neuropsychiatric studies. Those studies had not collected agonal factors nor had they corrected for unknown factors. This phenomenon especially exists in post-mortem brain samples. After we applied SVA to microarray data of samples from patients with SCZ, ASD and IBD, we found hypoxia-related pathways and oxygen level-related pathways. In the IBD data, which was not from brain tissue, we did not find any agonal-related pathways. This phenomenon suggests that the post-mortem brain is especially vulnerable to agonal factors. Moreover, we recommend a standardized data correction method to minimize the contribution of agonal factors. Normally, researchers can use linear regression analysis to correct for agonal factors similar to the measures used to correct for biological factors. If agonal factors were not documented, researchers can use SVA and PEER to adjust for unknown, unmodeled and latent sources of noise. These methods can detect variants induced by agonal factors. This is an important step for quality control in data preprocess. We also strongly suggest that researchers recheck previous results, since we identified several study results that were enriched in agonal-related modules and one study’s data confirmed to have been confounded by agonal factors. Our results provide a clear guidance for taking agonal factors into account and correcting for them in future research.

Our sample size was relatively small. We attribute the lack of significant DEGs or gene co-expression modules to these relatively small sample sizes, specifically, because of the limited number of *surgery* and *artificial respiration* phenotypes. A greater sample size will be necessary to validate the effects of unconsciousness, breathing difficulty, artificial respiration and premortem surgery. We also lacked a direct result replication for *fever-* and *infection-*related differentially expressed genes due to the widespread dearth of agonal factor recording in postmortem brain studies. For this reason we also lack a comprehensive record of agonal factors of gene expression data and are unable to evaluate overall agonal factors systematically. Furthermore, we validated only the co-expression network in different datasets using AFS instead of terminal states of fever or infection, which is not persuasive enough. Although we found a consistent cell-type enrichment module, larger datasets are still needed for validation.

## Methods

### Samples

Discovery data was collected from the ROSMAP project, including 263 samples with detailed information about each donor’s agonal conditions. Terminal state information included *fever, infection, surgery*, “unconc” for *unconsciousness*, “brthprb” for *difficulty breathing*, and “ventilat” for *artificial ventilation* within the hour (fever, infection, surgery, unconsciousness and difficulty breathing) or days (artificial ventilation) prior to death. The phenotypes *fever, infection* and *difficulty breathing* indicated that any of those experiences occurred within the three days prior to death; and, the phenotype of *surgery* indicated a major surgery with anesthesia in the two weeks prior to death. The sample size for each terminal state is shown in Extended Figure 1.

**Extended Figure 1.**
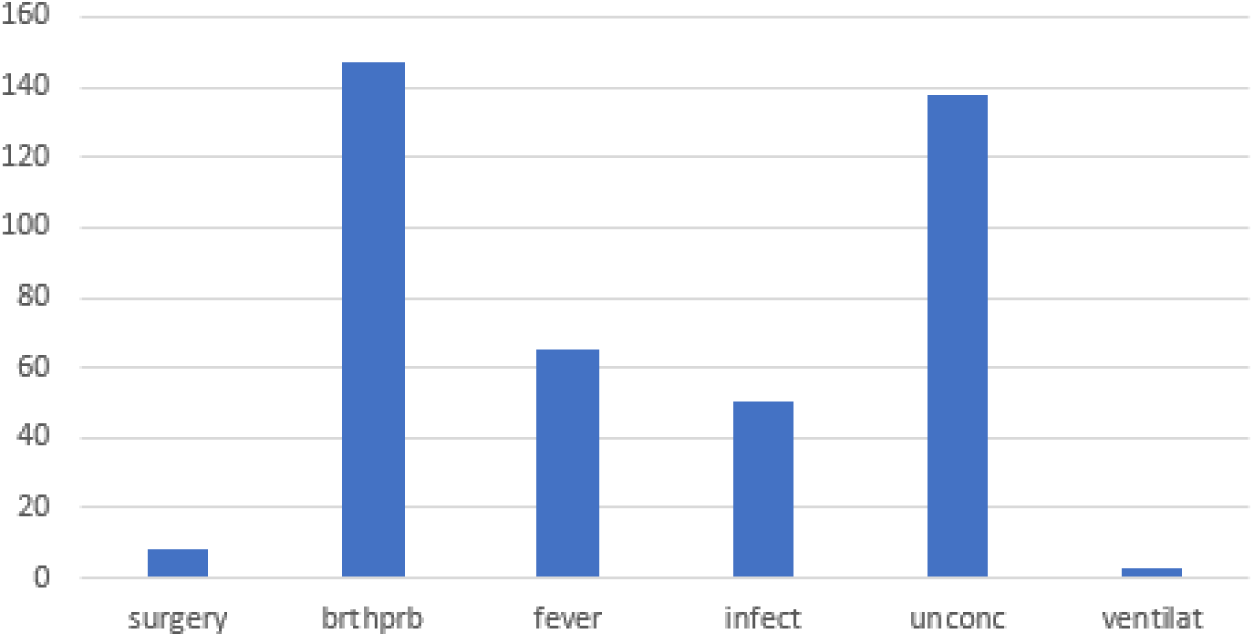
Sample size per terminal state.

Microarray data from Hagenauer’s group^9^ (GSE92538) was used for replication. We downloaded 335 samples in total with the agonal factor score (AFS), based upon the manner of death and terminal states combined. The manner of death is a categorical phenotype, ranging from 0 to 3, with 0 for a *violent, fast death*, 1 for *fast death of natural causes*, 2 for *intermediate death* and 3 for *slow death*. Terminal states include coma, medical condition (infection, sepsis), organ failure, head injury, hypoxia, brain death, mechanical respiration, and seizure. These are scored with 0 for “no” and 1 for “yes”. In these samples, 264 samples had AFS equal to 0, 45 had AFS equal to 1, 14 had AFS equal to 2, and 12 had AFS equal to 3. A detailed report on the agonal conditions of these samples was unavailable.

We also try to adjust latent variables of agonal factors in microarray data which did not documented manner of death or terminal states. We performed meta-analysis of different source of data (Extended table 1), including 5 microarray data of Schizophrenia, 3 microarray data of Autism Spectrum Disorder, and 2 microarray data of Inflammatory Bowel Disease (IBD). The source of microarray data of neuropsychiatric disorders is brain tissue, while the source of IBD is bowel tissue.

**Extended table 1.**
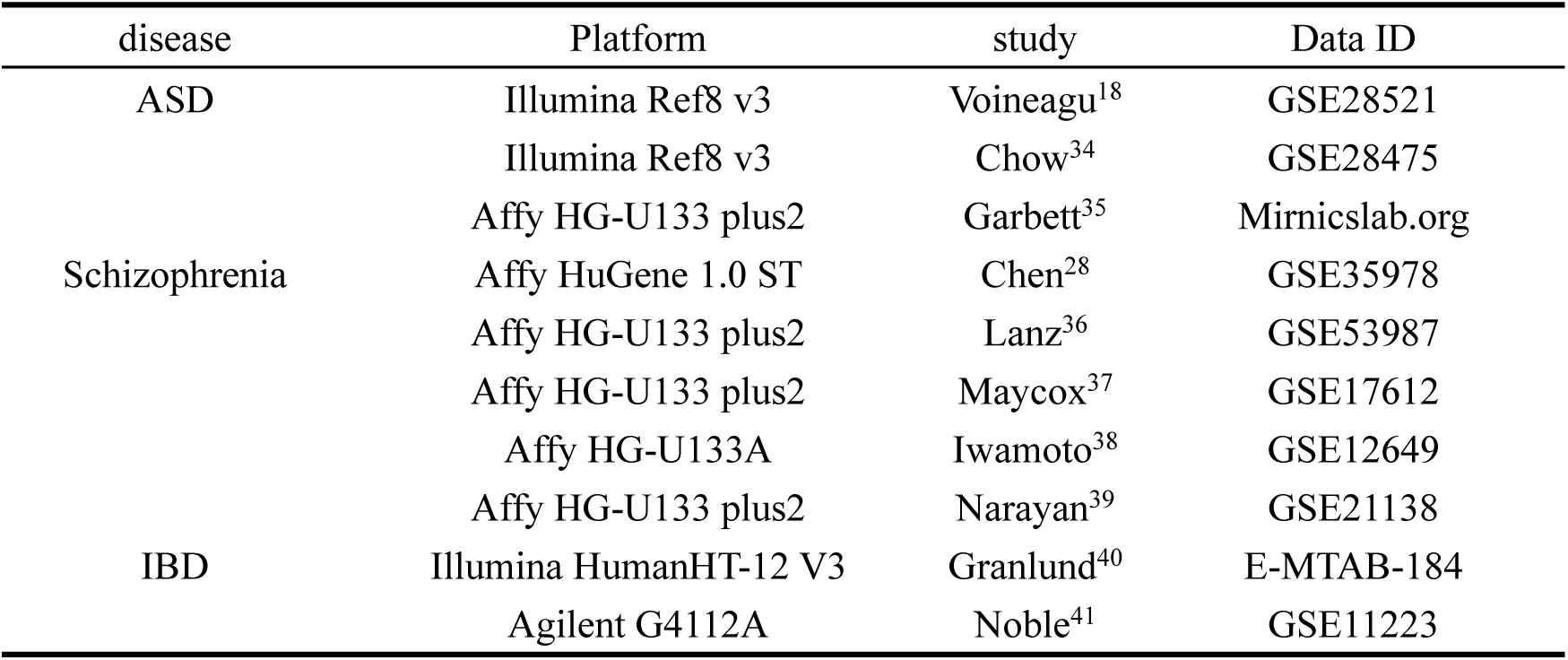
microarray data to adjust latent variables of agonal factors.

### Data preprocess

We performed quality control and data normalization. First, we performed Principle Component Analysis (PCA) and hierarchical clustering to filter outliers. We found sample ID 23690880 was potential outliers, so we removed this sample. Next, we filtered genes which were low expressed. The threshold of filtering is that expression value is less than 0.1 in 20% samples. We included 9092 genes in ROSMAP datasets and microarray data GSE92538 for following analysis. Then, we performed linear regression models corrected for the biological factors of the data. We corrected for factors including sex, racial group, Spanish ethnicity, years of education, age at death, postmortem interval, CERAD score (Consortium to Establish a Registry for Alzheimer’s Disease), Braak stage, and the National Institute on Aging (NIA)-Reagan Institute diagnosis criteria for Alzheimer’s disease. Finally, we used quantile normalization to remove technique variables.

### Statistical analysis

We performed differential gene expression using DESeq2, which transformed the data type into a gene expression matrix of integer values. In the discovery data, we used *fever, infection, unconsciousness, difficulty breathing*, and *artificial ventilation*, respectively, as the design matrix in DESeq2. Resulting P-values were corrected using the Benjamin-Hochberg (BH) procedure to control for multiple comparisons. We compared the differential gene expression effect size of the 5 agonal conditions using the pSI package and Pearson’s correlation. We performed all statistical analyses using R (v3.6.0).

### Cell deconvolution

We used cell deconvolution to estimate cell proportion of brain cell types in ROSMAP datasets. We used R package MuSiC^42^, which utilizes cell-type specific gene expression from single-cell RNA sequencing data to characterize cell type compositions from bulk tissue RNA-seq data. In our analysis, we used single-cell RNA sequencing data from ROSMAP^43^ as reference of cell deconvolution.

### Module construction and preservation testing

We identified the gene co-expression network using WGCNA. Before the analysis, we log transformed the data matrix to ensure normal distribution. We calculated a correlation matrix for all genes and chose the soft-threshold power of 12 to construct an approximate scale-free topology network. Networks were constructed using the blockwiseModules function. We chose the signed network type. The network dendrogram was created using an average linkage hierarchical clustering of the topological overlap dissimilarity matrix (1-TOM). Modules were defined as branches of the dendrogram using the hybrid dynamic tree-cutting method. Modules were summarized by their first principal component (ME, module eigengene) and modules with eigengene correlations of >0.9 were merged together. Modules were defined using Pearson correlation. Module (eigengene)-disease associations were evaluated using Pearson correlation. Significance values were FDR-corrected to account for multiple comparisons.

We used an additional dataset (GSE92538) to test the module preservation. Data preprocessing was the same as for the discovery data. We applied the Zsummary test to assess module preservation between expression datasets. The following thresholds are recommended: Zsummary < 2 implies no evidence for module preservation, Zsummary < 10 implies weak to moderate evidence, and Zsummary > 10 implies strong evidence for module preservation^44^.

### Cell-type enrichment analyses

Cell type enrichment was determined using the Zhang dataset, which uses cell type-specific expression datasets from human cortex brain samples from populations of neurons, astrocytes, oligodendrocytes, microglia and endothelial cells^45^. After normalizing and averaging replicate expression profiles for each cell type, a specificity index statistic (pSI) was calculated using the pSI package.

### Gene ontology enrichment

We performed Gene Ontology (GO) enrichment for biological process, molecular function and cellular components using the clusterProfiler v3.12.0 package in R^46^. The enrichment P-values were BH-corrected to control for multiple comparisons.

### Unknown factors detection

To detect unknown factors within the ROSMAP dataset, we employed SVA and PEER^47^ algorithms in R. The SVA package contains functions for identifying and building surrogate variables for gene expression data that could be used in subsequent analyses to adjust for unknown, unmodeled, or latent sources of noise. In SVA, we created a full model matrix, including race as a variable of interest. We detected 16 surrogate variables in total using the SVA package. PEER was used first to unearth patterns of common variation across the whole dataset and to create up to 15 assumed global hidden factors. Next, the correlation between terminal states and each of the 16 surrogate variables or 15 hidden factors was tested in the ROSMAP dataset. After that, factors showing a Pearson’s correlation test FDR-adjusted P-value smaller than 0.05 were included in linear regression analysis. The residual values from the regression analysis were used to correct the variables.

## Acknowledgements

We thank our patient donors and their families who have contributed to this research via postmortem brain tissue donations. We thank all members of the Pritzker Consortium (especially the University of California Irvine, Irvine, CA, United States of America) for producing the microarray data GSE92538. We thank Rush University and the ROSMAP project for the RNA-sequencing data. We also thank Richard F. Kopp and Liz Kuney from SUNY Upstate Medical University, for their language editing contribution, which greatly improved the manuscript.

## Funding

This work was supported by the Fundamental Research Funds for the Central Universities of Central South University (Grant No. 1053320184146), the National Natural Science Foundation of China (Grant Nos. 31970572, 31571312 and 81401114), the National Key R&D Project of China (Grant No. 2016YFC1306000), Innovation-driven Project of Central South University (Grant Nos. 2015CXS034 and 2018CX033), Hunan Provincial Natural Science Foundation of China (Grant No. 2019JJ40404) (to C. Chen), and the National Natural Science Foundation of China (Grant No. 31871276), the National Key R&D, Project of China (Grant No. 2017YFC0908701) (to C. Liu).

## Notes

### Competing Interest Statement

The authors have declared no competing interest.

